# A myofilament lattice model of *Drosophila* flight muscle sarcomeres based on multiscale morphometric analysis during development

**DOI:** 10.1101/2025.02.27.640547

**Authors:** Péter Görög, Tibor Novák, Tamás F. Polgár, Péter Bíró, Adél Gutheil, Csaba Kozma, Tamás Gajdos, Krisztina Tóth, Alexandra Tóth, Miklós Erdélyi, József Mihály, Szilárd Szikora

## Abstract

The indirect flight muscle is a widely used model for studying sarcomere structure and muscle development due to its extremely regular architecture. Nevertheless, precise measurement of the basic sarcomeric parameters remains a challenge even in this greatly ordered tissue. In this study, we identified several factors affecting measurement reliability and developed a software tool for precise, high-throughput measurement of sarcomere length and myofibril width. The accuracy of this new tool was validated against simulated images and blinded manual measurements. To extend the scope of this morphometric analysis to the sub-sarcomeric scale, we used electron and super-resolution microscopy to quantify myofilament number and filament length during myofibrillogenesis. These results provided novel insights into the dynamics of sarcomere growth, and enabled us to construct a refined model of sarcomere growth reaching to the level of individual myofilaments and providing a spatial framework for interpreting molecular localization data. These findings enhance our understanding of sarcomere assembly and offer a foundation for future studies of muscle development and function.

**SUMMARY STATEMENT:** This study identifies factors influencing sarcomere size measurements, introduces a validated tool for precise analysis, and presents a comprehensive model of *Drosophila* flight muscle sarcomere growth during muscle development.

## INTRODUCTION

The indirect flight muscles (IFMs) of *Drosophila* provide an excellent model for studying muscle organization and the molecular processes of myogenesis. The IFM is especially useful for investigating the assembly of myofilaments (thin and thick filaments) and sarcomeres, due to its regular structure, uniformity and synchronized growth during pupal development. Over the years, significant research, combined with advances in imaging techniques and the wide range of functional genomic tools available in *Drosophila* (Spletter et al., 2018, Sarov et al., 2016, Meiler et al., 2021, Zirin et al., 2020, Kanca et al., 2022, Loreau et al., 2023), has made the IFM one of the most well-studied muscle systems. Techniques like fluorescent nanoscopy can now reveal the detailed composition and arrangement of densely packed sarcomeric regions. Single-molecule localization microscopy, along with structural averaging, can pinpoint sarcomeric protein positions with precision under 10 nm, and has been used to reconstruct large sarcomeric protein complexes (Szikora et al., 2020a, González Morales et al., 2023, Schueder et al., 2023, Farkas et al., 2024). Additionally, *in situ* cryo-electron tomography and cryo-focused ion-beam scanning electron microscopy offer the possibility of detailed molecular and structural analysis of the myofilaments in their native state (Burbaum et al., 2021, Wang et al., 2021). The sarcomere lattice encapsulates the filamentary backbone of the sarcomeres, defining the number, length and arrangement of the thick and thin filaments, as well as the key sarcomeric regions like the H-zone, Z-disc, A-band and I-band. With the goal of achieving molecular reconstruction of IFM sarcomeres, the time is ripe to update existing morphometric data on the myofilament lattice in both mature and developing IFM sarcomeres, and to consolidate this information into a comprehensive model. Such models could improve our understanding of the mechanisms of sarcomere growth and provide geometric constraints necessary for interpreting data from fluorescent nanoscopy, enabling molecular modeling.

In this study, we provide a detailed analysis of IFM sarcomere growth. Building on previous work, we summarize the current knowledge on myofilament lattice morphometrics of IFM sarcomeres while addressing discrepancies in the literature. We also highlight the importance of measurement conditions and the need for precise reporting. Furthermore, we introduce a software tool that automates reliable size measurements from micrographs of IFM myofibrils. Finally, using a combination of conventional fluorescence microscopy, single-molecule localization microscopy and transmission electron microscopy (TEM), we present a model of the myofilament lattice of IFM sarcomeres, detailing the average number and length of the myofilaments, as well as the dimensions of the sarcomeric bands and zones during myofibrillogenesis.

## RESULTS AND DISCUSSION

### Discrepancies in reported IFM sarcomere morphometrics

Flight in *Drosophila* is driven by the coordinated action of the IFM, the largest muscle in flies, which is divided into Dorsal Longitudinal Muscles (DLMs) and Dorso-Ventral Muscles (DVMs) (Fig. 1A). The contractile machinery of the fibrillar IFM consists of uniform cylindrical myofibrils extending along the entire length of the muscle fibers. Each myofibril is composed of hundreds of serially connected, nearly uniform-sized sarcomeres (Fig. 1A) (Fernandes et al., 1991, Reedy and Beall, 1993). Consequently, sarcomere length and myofibril width (or diameter) are frequently reported morphometric traits.

**Figure 1.**
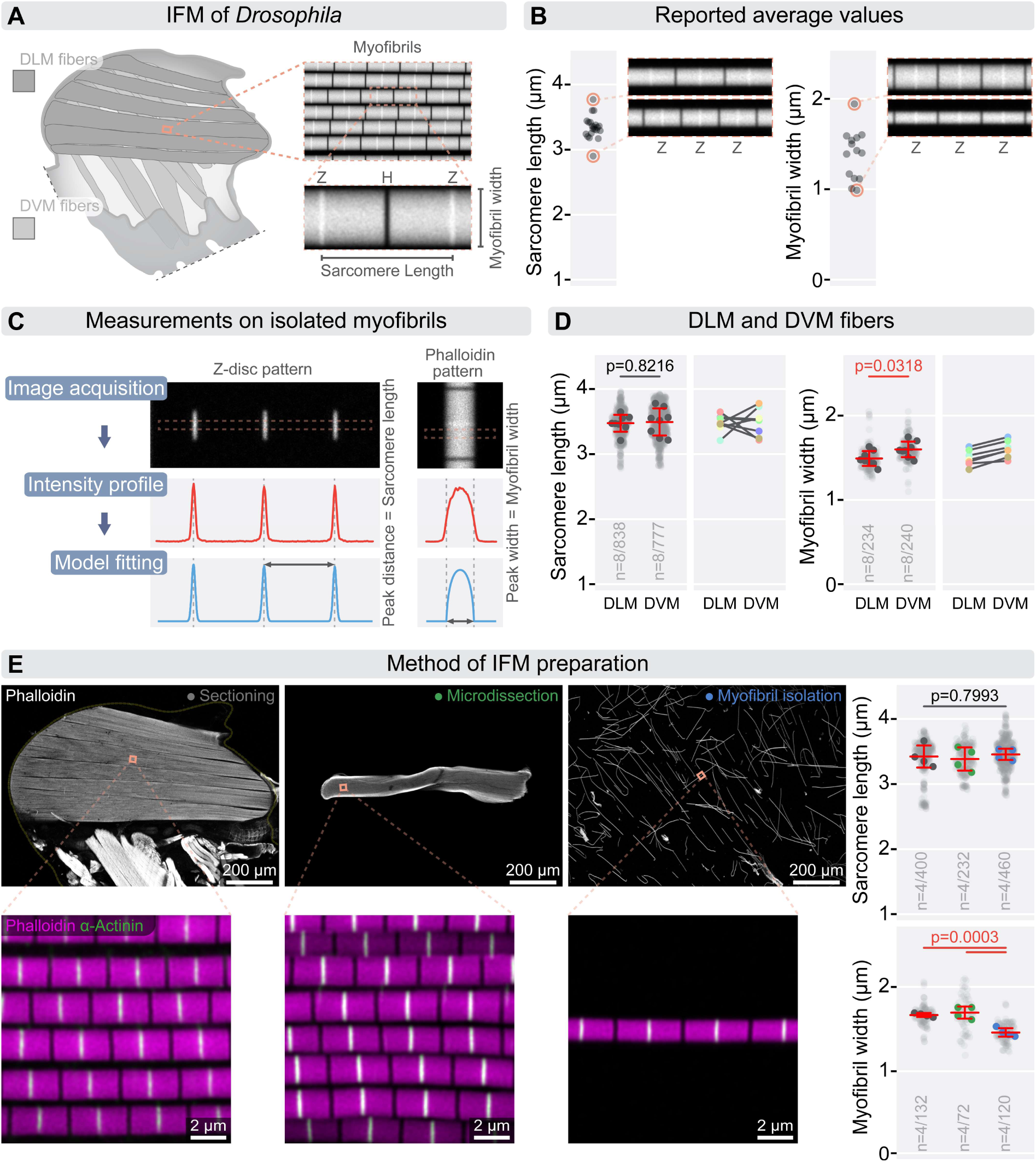
Morphometric measurements of IFM sarcomeres. **(A)** Schematic illustrating that indirect flight muscles (IFMs) are made up of the dorsal longitudinal muscles (DLM, shown in darker gray) and the Dorso-Ventral muscles (DVM, shown in lighter gray). The myofibrils consist of regular, uniform sarcomeres with characteristic lengths and diameters. The Z-discs (Z) at the borders of sarcomeres and the central H-zone (H) are labeled. **(B)** Plots displaying previously reported averages for sarcomere length and diameter, along with simulated images on the right that highlight the differences of the published measurements. **(C)** Panels outlining the process for measuring sarcomere length and diameter. Sarcomere length is determined from images where the Z-disc is labeled, while myofibril diameter is measured from phalloidin-labeled samples. After image acquisition, an intensity profile is recorded either along or perpendicular to the myofibril. This profile is then fitted with the appropriate Gaussian or Disc Model function to calculate sarcomere length and myofibril diameter, respectively. **(D)** A comparison of sarcomere length and diameter between DLM and DVM fibers reveals that while sarcomere lengths in these fibers are not significantly different (p=0.821), DVM fibers consistently exhibit larger diameters in every tested animal (p=0.031). Panels on the right show connected mean values for DLM and DVM fibers from the same animal. Statistical analysis was performed using an unpaired t-test. **(E)** Low- and high-resolution images illustrate commonly used methods for preparing IFM samples. F-actin is stained with phalloidin (gray or magenta), and Z-discs are labeled with α-Actinin (green). The impact of these preparation methods on sarcomere morphometrics is compared. While sarcomere lengths do not differ significantly between methods (p = 0.799), the diameter of individual myofibrils is significantly smaller in isolated myofibrils compared to sectioned or microdissected samples (p = 0.003). Statistical analysis was conducted using one-way ANOVA with Tukey’s multiple comparison test. Light gray dots represent individual measurements of sarcomere length and myofibril diameter, while the larger dots indicate the mean values from independent experiments. Error bars represent the mean and s.d. of independent experiments. “n” refers to the number of independent experiments and the number of replicates. Raw data used to generate the plots presented in this figure are available in the source data file (Fig1SourceData).

Despite the remarkable regularity of IFM sarcomeres, various laboratories tend to report significantly different values, with average sarcomere lengths ranging from 2.9 µm to 3.77 µm and myofibril widths from 0.98 µm to 1.94 µm in adult flies (Fig. 1B) (Mardahl-Dumesnil and Fowler, 2001, DeAguero et al., 2019, Spletter et al., 2018, Shwartz et al., 2016, Orfanos et al., 2015, Reedy and Beall, 1993, Fernandes and Schöck, 2014, Kooij et al., 2016, González Morales et al., 2023, González-Morales et al., 2019, Chakravorty et al., 2017, Katzemich et al., 2015, Perkins and Tanentzapf, 2014, Dhanyasi et al., 2020, Deng et al., 2021, Nongthomba et al., 2003, Cripps et al., 1999, Beall et al., 1989, Kao et al., 2021, Green et al., 2018, Miller et al., 2008, Nikonova et al., 2024, Tanner et al., 2011).

To address these discrepancies, we conducted a review of the published measurement methods. This analysis revealed several significant factors contributing to the variability in reported measurements.

### An automated method to accurately measure the sarcomeric parameters

First, we decided to assess the accuracy and precision of our routinely used measurement methods. Sarcomeres are bordered by Z-discs, and by definition, their length is equal to the distance between two adjacent Z-discs. Z-disc markers, such as α-Actinin or ZASPs, are typically used to label these discs, enabling relatively simple measurements from confocal micrographs (Fig. 1C). The other major parameter, myofibril diameter, is most often measured using phalloidin labeling, which can, however, be challenging sometimes because the edges of the myofibrils are not sharply defined, even against a uniform background (Fig. 1C).

For the analysis, we used isolated individual myofibrils because they are ideal for both localization studies and morphometric measurements. IFM myofibrils are easy to isolate and prepare, even for researchers with little experience, and produce highly reproducible results. Once isolated, the myofibrils lie flat on the coverslip, enabling clear, undistorted imaging with the highest possible resolution. Their orientation, parallel to the focal plane of the objective lens, allows for accurate assessment of two-dimensional projections without interference from surrounding structures. Additionally, morphometric analysis, using these images can be easily automated and these preparations also provide a random mixture of myofibrils, potentially from multiple animals, which strengthens the reliability of the analysis.

To determine the precision and accuracy of sarcomere length and myofibril diameter measurements, we generated artificial images of IFM myofibrils with known dimensions, simulating the image formation process (Fig. S1A). Since most previously reported data were obtained through manual measurements, we conducted a blinded test to evaluate their reliability. This test revealed significant variability between individuals, often considerably underestimating myofibril diameter (Fig. S1E).

Several automated tools for morphometric quantification of sarcomeres have been developed in recent years (Baheux Blin et al., 2024, Spletter et al., 2018, Stein et al., 2022, Neininger-Castro et al., 2023, Zhao et al., 2021, Morris et al., 2020, Toepfer et al., 2019, Pasqualin et al., 2016). While these tools generally provide accurate sarcomere length measurements, none can adequately measure myofibril diameter from regular side view images. Additionally, some tools require programming expertise or involve manual input for selection, which slows down the analysis process. To address these limitations, we developed a software tool called IMA, that automatically segments myofibrils from micrographs, applies spline fitting, and fits the appropriate models to intensity profiles to accurately calculate both sarcomere length and myofibril width (Fig. 1C, S2). For sarcomere length, the software analyzes intensity profiles along the myofibrils in the Z-disc marker channel, fitting them with multiple Gaussian functions to determine peak distances, which correspond to sarcomere size. For myofibril diameter, the software extracts intensity profiles perpendicular to the myofibrils from the phalloidin channel and fits them with a “disk function” to produce precise diameter estimates (Fig. 1C, S2) (additional details are provided in the IMA User Guide and the Methods section.).

The accuracy and precision of the tool were validated using simulated IFM images with known dimensions. In practical applications, this software not only improves measurement accuracy but also significantly reduces analysis time by 15 to 20 folds, hence subsequently, this new tool was applied for our individual myofibril measurements.

### IFM sarcomere morphometrics is affected by sex, age, fiber type and sample preparation

While reconsidering the literature, we noticed that the sex of the flies is often not specified, even though female pupae develop slightly faster than males, leading to earlier eclosion (Bainbridge and Bownes, 1981). When comparing newly eclosed males and females, we observed that males have slightly shorter sarcomeres and narrower myofibrils (Fig. S1C). This difference in sarcomere length disappears 12 hours after eclosion (AE), and the myofibril diameter difference resolves between 24 and 48 hours AE (Fig. S1C). These measurements highlight that factors such as the sex and age of the flies can affect measurement outcomes. Additionally, since the developmental rate of *Drosophila* is highly influenced by temperature, it is important to conduct experiments at a consistent temperature and ensure that comparisons are made under similar conditions.

The IFMs consist of two muscle groups: the DLMs, which develop using larval muscle templates, and the DVMs, which form *de novo* (Fernandes et al., 1991, Costello and Wyman, 1986). Despite their different developmental origin, the sarcomeres of these muscles are generally considered structurally identical. To test this, we microdissected DLM and DVM fibers from adult females (24 hours AE) and isolated individual myofibrils from them to compare their sizes. Our results showed that while sarcomere lengths were the same in both muscle groups, DVM myofibrils were consistently wider than those of the DLM (Fig. 1D), indicating that, even if slightly, muscle fiber type (DLM, DVM, or a mix) is another factor that can affect measurements.

Different studies often utilize flies with different genetic background. To determine whether these genetic variations influence IFM morphometric measurements, we tested several commonly used strains (*white^1118^, Oregon-R* and *Canton-S* often used as wild type controls, and *mef2-Gal4/+* as a frequently used muscle-specific Gal4 driver line), and found no significant differences in their sarcomeric parameters (Fig. S1F), indicating that the genetic background at this level has a negligible influence, and any of these strains can serve as an accurate control. Despite these findings, we trust that application of the best possible genetic matches will remain the most appropriate controls in future studies.

Sample preparation introduces numerous variables with a potential to alter the morphometrics of IFM sarcomeres. Common methods for preparing IFM for imaging include sectioning (via vibratome or cryosectioning), microdissection, or isolating individual myofibrils. The tissue is typically fixed using an aldehyde-based fixative, such as formaldehyde or glutaraldehyde. After immunostaining, the samples are embedded in either a liquid or hardening medium. While sarcomere length appears to be stable, the diameter of myofibrils is highly susceptible to changes during the various steps of sample preparation (Fig. 1E; Fig. S1 B, D). One of the most impactful steps is tissue embedding; specifically, using a hardening mounting medium significantly reduces myofibril diameter without affecting sarcomere length (Fig. S1B). This effect is less pronounced when myofibrils are fixed with glutaraldehyde (Fig. S1D) or when larger tissue sections are used (Fig. 1E). A previous study also highlighted that fixation before or after sectioning can also significantly affect the measured values (DeAguero et al., 2019). Although the hardening media causes shrinkage artifacts, we kept using them during our studies because they counteract the effects of lattice overhydration, which occurs when the sarcolemma is removed. Sarcolemma removal (or skinning) disrupts osmotic regulation, leading to overhydration of the myofilament lattice, increased lattice spacing, and thicker myofibrils (Miller et al., 2008, Tanner et al., 2012, Tanner et al., 2011). In contrast, dehydration—necessary for electron microscopy—reduces lattice spacing and shrinks myofibril diameter significantly (Chakravorty et al., 2017). Myofilament lattice spacing can be measured *in vivo* using small-angle X-ray scattering (Irving and Maughan, 2000, Miller et al., 2008, Tanner et al., 2012, Tanner et al., 2011), while the number of myofilaments can be determined from EM cross-sections (Fernandes and Schöck, 2014, Shwartz et al., 2016, Chakravorty et al., 2017). Together, these measurements allow for the estimation of *in vivo* myofibril diameter, which ranges from 1.54 to 1.68 µm. Our findings indicate that myofibrils prepared using hardening media have a diameter just within the range of these estimates (1.54 µm), unlike the diameter of those embedded in liquid media (2.2 µm), and for this reason we recommend using hardening media whenever possible.

Overall, it appears that sarcomere length and myofibril width are both affected by numerous factors in the commonly used experimental conditions. Curiously, however, sarcomere length is much less sensitive to these conditions than myofibril width, which varies in a broad range depending on the experimental circumstances and preparation method. These findings highlight the crucial importance of careful controlling and reporting of our methods to ensure the highest level of consistence in our IFM investigations.

### Synchronized growth and developmental phases of IFM myofibrillogenesis

During myofibrillogenesis, IFM sarcomeres undergo a substantial, synchronized growth in both length and diameter. Until now, only a few studies have provided detailed insights into this process (Reedy and Beall, 1993, Orfanos et al., 2015, Spletter et al., 2018). While these studies outlined a similar overall pattern of sarcomere growth (Fig. S3A), some ambiguities remained as to several questions: What is the exact size of the earliest detectable sarcomeres? Is sarcomere growth a continuous process, or does it include pauses? When do sarcomeres reach their final size?

Having our improved measurement methods in hand, we decided to revisit these questions and examine the developmental dynamics of IFM growth. To this end, individual myofibrils were isolated from twelve developmental time points (spanning from 36 hours after puparium formation (APF) to 96 hours AE) in six independent experiments using mixed-sex wild-type flies (Fig. 2B). We then measured sarcomere length and myofibril width (Fig. 2A). Our findings showed that immature, periodic myofibrils start forming around 36 hours APF at 25°C. We isolated short, intact myofibrils consisting of 4–12 sarcomeres from these young pupae, and found that their sizes were comparable to those measured in microdissected muscles, confirming their structural integrity (Fig. S3B). Although α-Actinin showed a periodic Z-disc pattern in microdissected immature myofibrils, this pattern was largely absent in pre-extracted individual myofibrils (Fig. S3B), suggesting that this protein is loosely associated with the immature Z-discs and can be easily disrupted by detergents. Stable association of α-Actinin with Z-discs appeared between 48 and 60 hours APF. Among the Z-disc markers we tested, only the Sls700 *Drosophila* Titin-specific B2 and Kettin Ig16 antibodies displayed stable, detergent-resistant Z-disc localization in young pupae (36–48 hours APF) (Fig. S3B, S4E), suggesting that the giant elastic proteins play a key role in early Z-disc and myofibril assembly.

**Figure 2.**
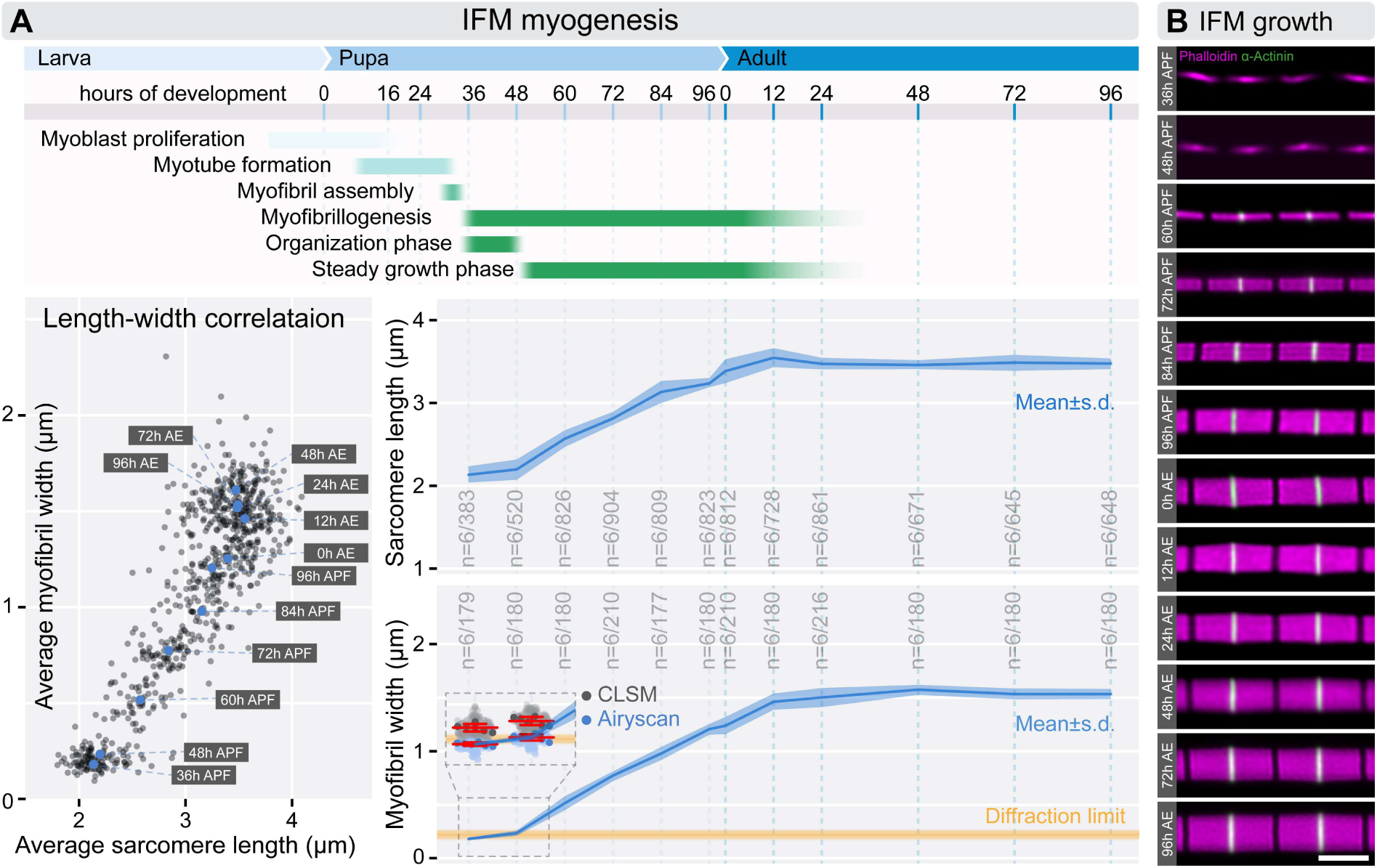
Growth of IFM sarcomeres during myofibrillogenesis. **(A)** The top panel provides a schematic outlining of the timeline and key steps of IFM myogenesis. In the bottom left, a plot illustrates the correlation between sarcomere length and myofibril width across myofibrillogenesis. Gray dots indicate mean values for individual myofibrils, while larger blue dots represent averaged means at specific time points. The plots on the right display the average sarcomere length and myofibril width measured across 12 time points, from 36 hours APF to 96 hours AE, following the timeline depicted in the schematic. “n” indicates the number of independent experiments and the number of replicates. A yellow line highlights the theoretical resolution limit of optical microscopy, with an inset showing that conventional light microscopy (LSM) overestimates the diameter of early myofibrils. In contrast, Airyscan imaging reveals that these diameters are below the diffraction limit. **(B)** Images of isolated individual myofibrils showcase the growth of IFM sarcomeres during myofibrillogenesis. F-actin (magenta) is stained with phalloidin, and Z-discs (green) are labeled with α-Actinin. Scale bar: 2 µm. Raw data used to generate the plots presented in this figure are available in the source data file (Fig2SourceData).

While measuring the diameter of early myofibrils (36 and 48 hours APF), we noticed an apparent lower limit which matched the theoretical diffraction limit of conventional fluorescent microscopes (Fig. 2A). To overcome this limitation, we used Airyscan imaging and found that myofibrils are considerably thinner than previously assessed by regular confocal microscopy (Spletter et al., 2018), though still thicker than measured on EM sections (Reedy and Beall, 1993) (Fig. 2A).

Between 36 and 48 hours APF we only detected a modest increase in sarcomere size, but after this period a marked growth took place without significant changes in the assembly rate up to 96 hours APF (Fig. 2). Subsequently, we observed a sharp increase in sarcomere length between 96 hours APF and the time of eclosion (0 hours AE), likely due to pre-stretching of IFM myofibrils as the thoracic exoskeleton expands. Our data showed that IFM sarcomere length and diameter generally peak and/or plateau in young adult flies between 12 to 24 hours AE (Fig. 2A). Interestingly, although the causal connection is unclear, we note that it precisely coincides with the isoform switch of Tropomodulin (Tmod), when the ∼43 kDa isoform responsible for thin filament elongation is replaced by a larger ∼45 kDa isoform (Mardahl-Dumesnil and Fowler, 2001). By 48 hours AE, any sex differences in myofibril diameter were equalized (Fig. S1C).

Based on these growth patterns, we identified two distinct phases of IFM myofibrillogenesis: the organization phase and the steady growth phase (Fig. 2A). During the organization phase, sarcomere size remained nearly constant, as observed previously (Kolley et al., 2024), while the number of sarcomeres increased significantly—from approximately 100 at 36 hours APF to about 230 at 48 hours APF (Spletter et al., 2018). After 48 hours APF, only a few additional sarcomeres were added, resulting in 260–270 sarcomeres per dorsal longitudinal muscle (DLM) myofibril by 60 hours APF (Spletter et al., 2018). In the steady growth phase, which spans from 48 hours APF to 24 hours AE, sarcomeres continued to elongate, and myofibril diameter expanded until both reached their mature size in adult flies between 12 to 24 hours AE (Fig. 2A).

### Assessing myofilament assembly and elongation during myofibrillogenesis

The growth of sarcomeres in both length and width involves two processes: elongation of the existing myofilaments (thin and thick filaments) and addition of new myofilaments to the sarcomere lattice. Studies, including ours, indicate that thin filaments extend specifically at their pointed ends, while thick filaments grow by adding myosin molecules at their ends. Additionally, newly formed myofilaments integrate at the periphery of the myofibrils, gradually expanding the structure in radial direction (Fig S4A) (Mardahl-Dumesnil and Fowler, 2001, Shwartz et al., 2016).

While standard fluorescent microscopy is sufficient to measure sarcomere length and diameter, determining the precise length and number of the myofilaments requires higher-resolution imaging. For this, we utilized fluorescent nanoscopy to measure myofilament length and electron microscopy to count myofilaments across four developmental stages (from 36 hours APF to 24 hours AE). To examine myofilament number and arrangement during myofibrillogenesis, we analyzed cross-sections of DLM myofibrils using transmission electron microscopy (Fig3. A, A’). At 36 hours APF, myofibrils appeared as disorganized clusters of filaments, with an average of 23 thick filaments per myofibril. By 48 hours APF, myofibrils began to adopt a more structured arrangement, establishing a lattice spacing typical of mature myofibrils (Fig. S5A). The number of thick filaments increased to an average of 32, forming a nearly perfect hexagonal pattern with a 1:3 ratio of thick to thin filaments, even though irregularities were still observed at the edges. At 72 hours APF, the lattice structure became fully organized, resembling the adult form, with an average of 134 thick filaments. In adult flies (24 hours AE), the number of thick filaments per sarcomere averaged around 846, consistent with previous research (Fernandes and Schöck, 2014, Shwartz et al., 2016, Chakravorty et al., 2017).

Exploiting that thin filaments are organized such that their pointed ends face the edge of the H-zone, while their barbed ends overlap at the Z-disc, their length can be measured by visualizing capping proteins like CapZ (a barbed end capper) and Tmod (a pointed end capper). This approach is effective from the later pupal stages (72 hours APF) till the young adults (24 hours AE; Szikora et al., 2020), where the Tmod and CapZ (i. e. cpa) signals form well-defined lines (Fig. S4B). However, in the IFM of early pupae (36 and 48 hours APF), both signals are weak and disordered, making such measurements impractical. Instead we used phalloidin-labeled myofibrils and visualized them with dSTORM microscopy to measure the length of the thin filament arrays (Fig. 3B). Phalloidin specifically binds to F-actin and, due to its small size, provides relatively uniform labeling across the entire myofibril without introducing ‘linkage errors’. Thin filament length could be deduced from sarcomere length, the width of the H-zone and of the Z-disc overlap (Fig. 3B, B’,B’’).

**Figure 3.**
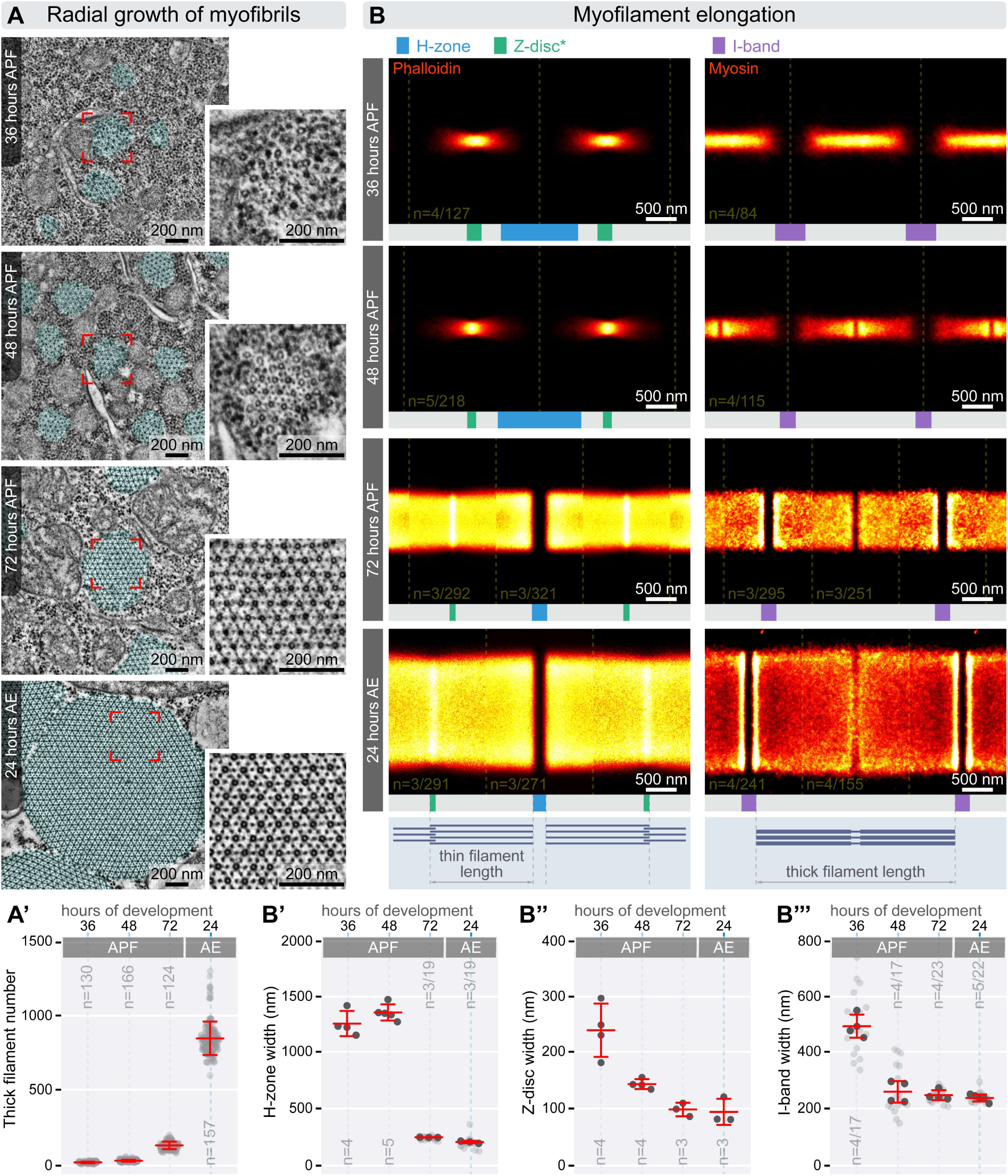
Characterization of myofilament number and length during myofibrillogenesis. **(A)** TEM cross-sections of DLM fibers highlight the radial growth of myofibrils (highlighted in cyan) throughout myofibrillogenesis. Higher-resolution insets on the right showcase changes in the lattice organization of myofilaments. **(A’)** The accompanying plot displays the average number of thick filaments observed in the TEM cross-sections, with the mean and s.d. indicated. **(B)** Averaged dSTORM reconstructions illustrate the growth and spatial organization of thin filaments (stained with phalloidin, on the left) and thick filaments (labeled with myosin antibodies, on the right) during myofibrillogenesis. **(B’-B’’’)** Plots showing measurements of H-zone width (B’), Z-disc width (B”) and I-band width (B’’’) derived from dSTORM images. Light gray dots represent the mean values for individual myofibrils, while dark gray dots indicate the mean values from independent experiments. The mean and s.d. of these experiments are provided. Raw data used to generate the plots presented in this figure are available in the source data file (Fig3SourceData).

At 36 hours APF, thin filaments were mostly misaligned, with the majority measuring less than 560 nm in length (with quite high standard deviations). The overlap of the thin filaments at the Z-disc was approximately 240 nm, consistent with the width of the electron-dense Z-bodies observed in longitudinal EM sections of myofibrils at this stage (Fig. S5B). While D-Titin (labeled with Sls700 B2 and Kettin Ig16) was already arranged in a bipolar pattern, the distribution appeared rather diffused. Additionally, the Z-discs lacked stable binding of α-Actinin and Zasp52 (Fig. S4C,E). By 48 hours APF we measured a thin filament length of ∼490 nm, which is somewhat shorter than at 36 hours APF. Nevertheless, we think it very unlikely that the actin filaments indeed shrink, instead we suspect that by 48 hours APF the sarcomeres attain a higher structural regularity than at 36 hours APF, without significant changes in filament length. Consistent with this idea, by 48 hours APF the Z-discs became more structured, with a reduced thin filament overlap of ∼140 nm, resembling the mature configuration, as also seen in longitudinal EM sections (Fig. S5B). The D-Titin markers also appeared more compact, likely indicating that filaments were aligning in register. However, stable associations of α-Actinin and Zasp52 were still absent (Fig. S4C,E). At 72 hours APF the thin filament array adopted a structure closely resembling its mature form, now with stable α-Actinin and Zasp52 associations (Fig. S4C). Thin filaments extended to ∼1330 nm, and the overlap at the Z-disc was reduced to ∼98 nm, which is only slightly larger than the ∼94 nm observed in mature sarcomeres. Longitudinal EM sections further supported these findings (Fig. S5B), and the Z-disc overlap size was consistent with measurements from the IFM of other species obtained through cryo-EM reconstructions (Yeganeh et al., 2023, Rusu et al., 2017). Finally, by 24 hours AE, thin filaments reached their final length of 1680 nm, and all Z-disc markers exhibited their fully matured, double-line pattern (Fig. S4C) (Szikora et al., 2020a).

To track the process of thick filament elongation we used monoclonal antibodies targeting the myosin heavy chain (Mhc). During the early pupal stages (36h and 48h APF) we used the 3E8 antibody, while for the later stages (72 hours APF and 24 hours AE) we applied the MAC 147 antibody. The 3E8 antibody labelled the isolated thick filaments uniformly (with the exception of the bare zone), suggesting that it recognizes an epitope in the S1 or S2 region (Fig. S4D). Similarly, the MAC 147 antibody recognizes an epitope in the S2 region of Mhc (Qiu et al., 2005). While the 3E8 antibody effectively labelled the thick filament arrays in the early pupal stages, the 3E8 signal significantly declined during the later stages (Figs. 3B, S4D). In contrast to this, the MAC 147 antibody labelled the A-band during the later stages with a distinct bare zone and with a strong intensity at the I-band/A-band border, but it was not suitable to label the myofibrils in the young pupae. The length of the thick filaments was calculated based on the average sarcomere length and the I-band width measured on the nanoscopic reconstructions (Fig. 3B,B’’).

At 36 hours APF, the thick filament array appeared fairly well-defined but it seemed somewhat misaligned since no bare zone was visible at the M-line. The array measured about 1640 nm in length on average, although individual thick filaments were probably slightly shorter. At this stage, Obscurin was only weakly detected using an antibody against its central domains (Ig14-16) (Fig. S4F). By 48 hours APF, the average thick filament length increased to ∼1940 nm, and a distinct bare zone became visible at the M-line, suggesting that the filaments were aligning in register. The I-band width at this point already resembled that of mature sarcomeres (Fig. 3B, B’’’). Additionally, the Obscurin Ig14-16 signal was now clearly seen as a single band along the M-line, as in previous reports for the adult muscles (Szikora et al., 2020a) (Fig. S4F). At 72 hours APF, both the width of the thick filament array and the length of the individual filaments had grown significantly, reaching approximately 2570 nm (Fig. 3B, B’’’). Finally, by 24 hours AE, the thick filaments attained their final length of ∼3240 nm, consistent with earlier studies (Katzemich et al., 2012, Gasek et al., 2016). At this stage, the Obscurin pattern became more compact along the longitudinal axis of sarcomeres, corresponding to a narrowed H-zone (Figs. 3B, B’; S4F).

Thus, the application of two high resolution microscopy methods (TEM and dSTORM) allowed us to collect single filament level information about the precise number and length of the myofilaments in the developing IFM. These data, combined with entire sarcomere level measurements, provided us with all knowledge to accurately determine the dimensions of every sarcomeric region, and served as an excellent starting ground for reconstruction of the IFM myofilament lattice.

### Myofilament lattice model of developing IFM sarcomeres

Using morphometric measurements from confocal, TEM and dSTORM microscopies, we generated sarcomere blueprints that represent the average sarcomere structure at specific developmental stages (see example in Fig. S6A). In the *Drosophila* IFM, myofilaments form a regular hexagonal lattice, maintaining a 3:1 ratio of thick to thin filaments. These filaments are arranged in a MyAc layer, with thin filaments consistently positioned at the midpoint between two thick filaments (Leonard and Bullard, 2006). Incorporating these structural constraints, we developed simplified models of average sarcomeres in cross-sectional and longitudinal views (see example in Fig. S6B). From these two-dimensional representations, we constructed three-dimensional models (Fig 4.; Supplementary Movie 1-4), assuming a hypothetical P321 symmetry for thin filaments in the Z-discs, similar to what is observed in the IFM of other insect species (Cheng and Deatherage, 1989, Yeganeh et al., 2023, Rusu et al., 2017). These models represent our current understanding of how the myofilament arrays in IFM sarcomeres expand in length and diameter during myofibrillogenesis.

**Figure 4.**
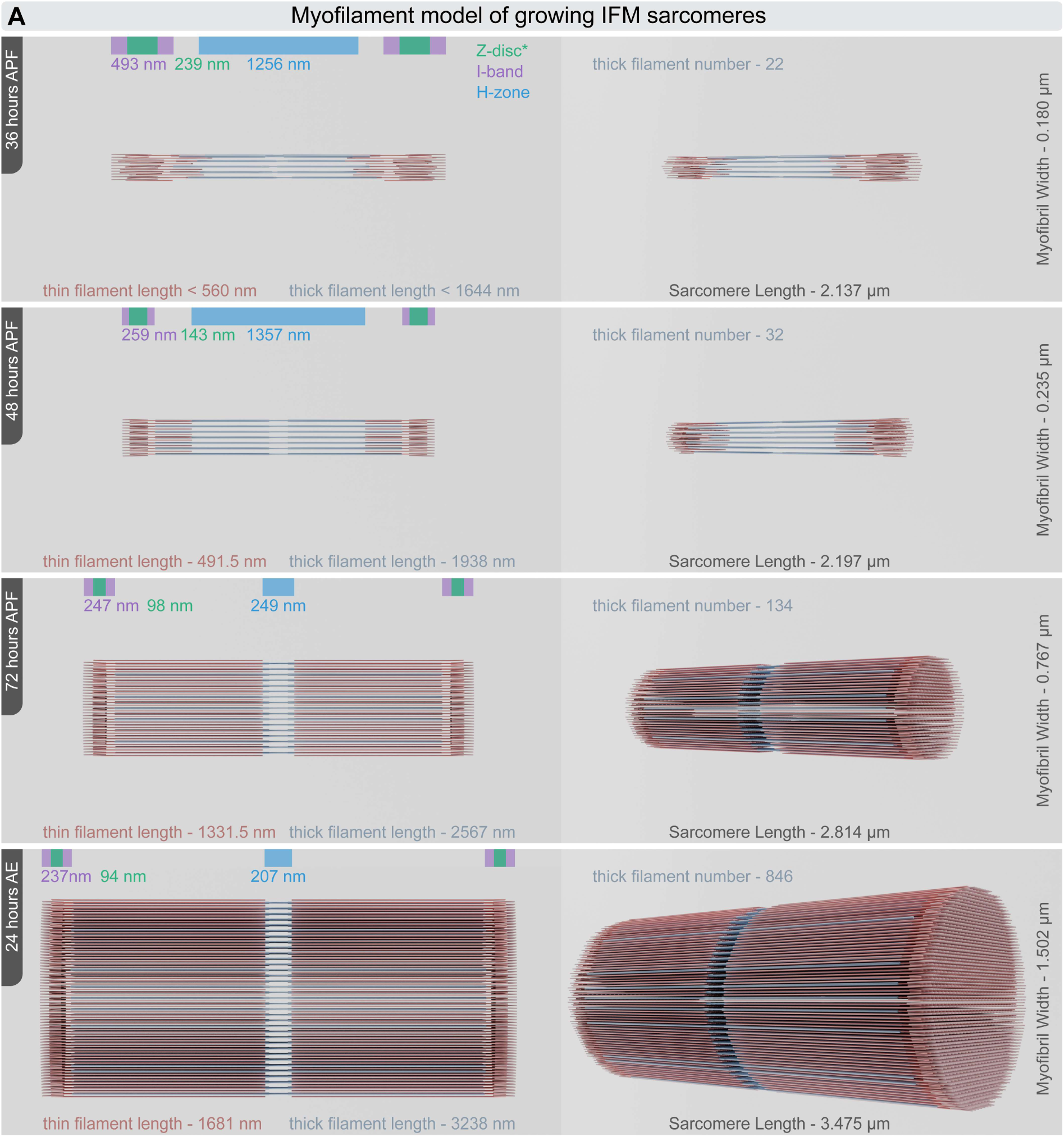
Myofilement model of developing IFM sarcomeres. **(A)** Three-dimensional models of sarcomeres illustrate the average number, size and arrangement of thick filaments (blue) and thin filaments (red) during myofibrillogenesis. The models on the left present orthographic views, while those on the right display rotated perspective views of the sarcomeres. Data presented in this figure are available in a table form in the source data file (Fig4SourceData).

## SUMMARY

The IFM is a broadly used model for investigating the molecular mechanisms underlying sarcomere structure and muscle development due to its highly regular, uniform and easily observable architecture. While this structural uniformity is not in question, we noted that reported measurements often exhibit significant variability as to the basic sarcomeric parameters. In this study, we identified several factors influencing these measurements and addressed them systematically. We found that a large portion of the differences can be attributed to the inaccuracy of the measurement methods. In addition, we revealed that although sarcomere length is consistent across sexes, muscle fiber types and experimental conditions, myofibril width is highly sensitive to the experimental methods, and it even exhibits a slight difference between DLM and DVM fibers.

To achieve precise and high-throughput measurements, we developed a software tool that segments myofibrils from micrographs, applies spline fitting, and fits the appropriate models to intensity profiles to accurately calculate sarcomere length and myofibril width. The accuracy of this new tool was validated using simulated IFM images with known dimensions, while its precision was confirmed by comparisons to manual measurements on blinded datasets. Transmission electron microscopy was applied to directly measure the number of myofilaments in IFM myofibril cross-sections during myofibrillogenesis. Additionally, dSTORM microscopy was used to determine the length of thin and thick filaments by measuring the width of the H-zone, I-band and Z-disc. These measurements were integrated into a comprehensive model describing sarcomere growth at the level of individual myofilaments. This model serves two primary purposes:

1. **Providing a spatial reference framework:** While fluorescent super-resolution microscopy precisely localizes sarcomeric molecules, interpreting these data can be challenging without a broader context of sarcomere ultrastructure. Our model offers a spatial framework that leverages the uniformity, regularity and symmetry axes of the sarcomere to accurately position specific molecules within the myofilament array (Szikora et al., 2020).
2. **Quantifying sarcomere growth dynamics:** The single filament level analysis during IFM development provided new information as to the dynamics of the growth process. During the first phase, from 36 to 48 hours APF, incorporation of a single thick filament (or six thin filaments) requires approximately 65 minutes. Between 48 and 72 hours APF, this process accelerates reducing the integration time to around 15 minutes, and from 72 hours APF to 12 hours AE, the incorporation time further decreases to about 4.5 minutes, eventually slowing down when the sarcomeres attain their mature size. These data can now be used as a cornerstone, or at least a reference point for comparisons of the developmental dynamics of IFM growth in different conditions.

All in one, we believe that this model could provide a foundation for future studies exploring the molecular mechanisms of myofilament elongation and formation. Moreover, it holds a great potential for a wide range of applications in understanding sarcomere dynamics at the molecular level.

## MATERIALS AND METHODS

### Drosophila Stocks

Flies were raised on cornmeal medium and maintained at 25°C under standard laboratory conditions. Morphometric analysis of sarcomere growth was conducted using the *w^1118^* strain. Stocks of *Oregon-R*, *Canton-S*, and *mef2-Gal4* (BDSC #27390), (Ranganayakulu et al., 1996) were used as controls. For staging, white pre-pupae or freshly hatched flies were collected and maintained until they reached the desired age. In the actin incorporation assay, the *UAS-Act88F::GFP* transgene (BDSC #9253) (Röper et al., 2005) was expressed using either the *duf-Gal4* (Menon and Chia, 2001) or *fln-Gal4* (Bryantsev et al., 2012) driver.

### Sample Preparation for Fluorescent Microscopy

#### IFM Agarose Sectioning

Thoraces were separated from the heads and abdomens using fine forceps, with the heads and abdomens discarded. The isolated thoraces were immediately placed in 4% paraformaldehyde (Thermo Fisher Scientific; paraformaldehyde, 16%) in relaxing solution (100 mM NaCl, 20 mM NaPi at pH 7.0, 5 mM MgCl2, 5 mM EGTA, and 5 mM ATP) and fixed overnight at 4°C. After a brief rinse in PBS, the thoraces were submerged in warm 5% agarose ((Lonza, SeaKem LE Agarose) prepared in PBS) and positioned appropriately. Once the agarose solidified, the blocks were glued to a sample holder and inserted into a vibratome (Microm HM 650 V). The lateral side of the embedded thorax was oriented toward the vibratome blade, which was submerged in PBS. Using the vibratome, 120 µm thick sections were collected and stored in PBS at 4°C. Standard immunohistochemistry procedures were then applied to the sections.

#### IFM Microdissection

Thoraces were separated from the heads and abdomens using fine forceps, discarding the heads and abdomens. The thoraces were then bisected along the sagittal midline and immediately fixed in 4% paraformaldehyde (Thermo Fisher Scientific; paraformaldehyde, 16%) prepared in relaxing solution for 20 minutes on ice. After fixation, IFM muscle fibers were carefully detached from the hemithoraces, transferred to PBS, and stored at 4°C. Standard immunohistochemistry procedures were subsequently applied to the isolated fibers.

#### IFM Individual Myofibril Preparation

Individual myofibrils were isolated from the IFM using a modified protocol based on Burkart et al., 2007. Briefly, bisected hemithoraces were incubated on ice in relaxing solution containing 50% glycerol for 2 hours. The DLM and/or DVM fibers were then carefully dissected from the hemithoraces and dissociated in an Eppendorf tube by pipetting with 0.5% Triton X-100. The dissociated myofibrils were centrifuged at 10,000 × g for 2 minutes at 4°C, and the pellet was re-suspended in 200 µl of 1× relaxing solution by pipetting. This centrifugation and washing process was repeated twice. After the final centrifugation, the fibers were re-suspended in relaxing solution, and 20 µl of the suspension was placed on a glass coverslip and fixed for 20 minutes either with 4% paraformaldehyde (Thermo Fisher Scientific; paraformaldehyde, 16%) or with 2% glutaraldehyde (Electron Microscopy Sciences; Glutaraldehyde, 8%) in relaxing solution. See (Szikora et al., 2020b) for more details.

#### Thick Filament Isolation

Thick filaments were isolated from the IFM following the protocol outlined in Katzemich et al., 2012. In summary, individual myofibrils (prepared as described above) were subjected to mild Calpain digestion to separate the thick filaments from the Z-disc (Contompasis et al., 2010, Reedy et al., 1981). The myofibrils were resuspended in 30 µl of a solution containing 20 mM imidazole (pH 6.8), 1 mM EDTA, 1 mM EGTA, 5 mM β-mercaptoethanol, and 30% glycerol, along with 15 mM Calpain-1 (Merck). CaCl_2_ was added to achieve a final concentration of 2 mM, and the mixture was incubated at room temperature for 30 minutes. To halt the digestion, 100 µl of relaxing solution and 100 mM Calpain Inhibitor I (Merck) were added. The suspension was then passed through a 20G needle five times to separate the filaments and centrifuged at 1500 × g for 3 minutes to remove undigested myofibrils. The resulting suspension was applied to a glass coverslip and fixed for 20 minutes with 4% paraformaldehyde (Thermo Fisher Scientific; paraformaldehyde, 16%). Isolated and fixed thick filament samples were then subjected to standard immunohistochemistry procedures.

### Immunohistochemistry

Fixed tissues were washed three times and blocked in PBS-BT for 2 hours at room temperature (30 minutes for individual myofibril samples). The following primary antibodies were applied in blocking solution and incubated overnight at 4°C:

α-Actinin (DSHB 2G3-3D7, mouse, 1:100)

Cpa (gift from Florence Janody, rabbit, 1:100; Amândio et al., 2014)

GFP (Abcam ab13970, chicken, 1:1000)

Kettin Ig16 (DSHB BB17/29.5, rat, 1:200)

Myosin 3E8 (DSHB 3E8-3D3, mouse, 1:1000)

Myosin MAC147 (DSHB BB7/21.14, rat, 1:1000)

Obscurin Ig14-16 (gift from Belinda Bullard, rabbit, 1:400; Burkart et al., 2007)

Sls700 B2 (gift from Belinda Bullard, rabbit, 1:200; Lakey et al., 1990)

Tmod (gift from Velia Fowler, rat, 1:200)

Zaps52 (DSHB 1D3-3E4, mouse, 1:400)

After additional washes, secondary antibodies were applied for 2 hours at room temperature. Detection was carried out with highly cross-absorbed goat anti-rabbit, anti-mouse, anti-rat or anti-chicken IgG secondary antibodies conjugated to Alexa Fluor 405, Alexa Fluor 488, Alexa Fluor 546 or Alexa Fluor 647 (Life Technologies, 1:600). F-actin was labeled with Alexa Fluor 488-, Alexa Fluor 546- or Alexa Fluor 647-phalloidin (Life Technologies, diluted in methanol, 1:200 or 1:800 for dSTORM imaging). After thorough washes, the samples were mounted in antifade reagent (ProLong Gold, P36930; Life Technologies), glycerol (90% glycerol, 10% PBS) or stored in PBS until dSTORM imaging.

### Confocal Laser Scanning Microscopy

Confocal images were acquired using a Zeiss LSM800 Airyscan microscope with either a Plan-Apochromat 63×/1.40 Oil DIC M27 or an EC-Plan-Neofluar 10×/0.30 M27 objective lens. Images were captured at the Nyquist rate to ensure optimal resolution, utilizing the full dynamic range of the GaAsP or Airyscan detectors.

### Super-Resolution dSTORM Imaging

Super-resolution imaging was conducted as previously described (Szikora et al., 2020a, Szikora et al., 2020b). Imaging was performed using a custom-built inverted microscope system based on a Nikon Eclipse Ti-E frame, equipped with a Nikon CFI Apo 100×, NA = 1.49 objective lens. Excitation was provided by a 647 nm laser (MPB Communications Inc., Pmax = 300 mW), with the intensity adjusted to 2–4 kW/cm² at the sample plane via an acousto-optic tunable filter (AOTF). A 405 nm laser (Nichia, Pmax = 60 mW) was used for molecule reactivation. Images were captured with an Andor iXon3 897 BV EMCCD camera (512 × 512 pixels, 16 μm pixel size) as frame stacks for dSTORM imaging at reduced image size. A fluorescence filter set (Semrock, LF405/488/561/635-A-000) and an additional emission filter (AHF, 690/70 H Bandpass) were used to separate excitation and emission light. During imaging, the perfect focus system of the microscope maintained sample focus with precision <30 nm. Before imaging the storage buffer was replaced with a GLOX switching buffer (van de Linde et al., 2011), and the sample was mounted onto a microscope slide. Typically, 20,000–50,000 frames were captured with an exposure time of 20–30 ms. Image stacks were analyzed using rainSTORM localization software (Rees et al., 2013), where single-molecule images were fitted with a Gaussian point spread function, and the center positions of fluorescent molecules were determined. Localization data were filtered based on intensity, precision (<20 nm), and standard deviation (0.8 ≤ σ ≤ 1.0). Drift caused by mechanical or thermal effects was corrected using a correlation-based blind drift correction algorithm. See (Szikora et al., 2020b) for more details.

### Transmission Electron Microscopy

Dissected hemithoraces were fixed overnight at 4°C in a solution containing 3.2% paraformaldehyde, 0.5% glutaraldehyde, 1% sucrose, and 0.028% CaCl₂ in 0.1 N sodium cacodylate buffer (pH 7.4). The samples were then washed twice in 0.1 N sodium cacodylate buffer (pH 7.4) overnight at 4°C, washed for 15 minutes in distilled water, and post-fixed for 1 hour in 1% osmium tetroxide (Sigma-Aldrich) in distilled water. After osmium fixation, the samples were rinsed in distilled water for 10 minutes and dehydrated in a graded ethanol series (50% to 100%) for 10 minutes at each concentration (twice in 100%). Following dehydration, the muscles were treated with propylene oxide for 5 minutes (Molar Chemicals) and embedded in an epoxy-based resin (Durcupan ACM; Sigma-Aldrich). Mixture of propylene oxide and epoxy-based resin (3:1, 1:1, 1:3) were added to the samples for 1-1-1 hours. After, samples were processed through pure resin twice for 1 hour, and through pure resin again in room temperature overnight. The resin blocks were polymerized at 56°C for 48 hours. Once polymerized, the blocks were etched, and 50 nm ultrathin sections were cut using a Leica Ultracut UCT ultramicrotome. The sections were mounted on single-hole, formvar-coated copper grids (Electron Microscopy Sciences) and stained to enhance contrast. Staining was performed using 2% uranyl acetate in 50% ethanol and 2% lead citrate in distilled water (both from Electron Microscopy Sciences). The ultrathin sections were examined using a JEM-1400Flash transmission electron microscope (JEOL). Muscle cross-sections were systematically screened at low magnifications (500–2000×), and images were captured at higher magnifications - 12,000× for cross-sections and 8000× for longitudinal sections. Images were recorded as 16-bit grayscale files using a 2k×2k high-sensitivity scientific complementary metal-oxide-semiconductor (sCMOS) camera (Matataki Flash, JEOL) and saved in tagged image file format.

### Image Processing and Analysis

#### Confocal images

Sarcomere length and diameter were measured directly from raw .czi files using a custom algorithm called *Individual Myofibril Analyser* (IMA). Isolated myofibrils were analyzed in Automatic Mode, while intensity profiles were manually selected for microdissected or sectioned muscle samples. The source code and a detailed user guide are available online (https://github.com/GorogPeter94/Individual-Myofibril-Analyser-IMA-/tree/main). The presented images were restored using Huygens Professional Software 23.10 (Scientific Volume Imaging). Optical sections were displayed as average projections, with brightness and contrast linearly adjusted in Fiji (Schindelin et al., 2012).

#### TEM images

For presentation purposes, the contrast of TEM images was enhanced using the Enhance Local Contrast (CLAHE) method in Fiji (Schindelin et al., 2012). Myofilament numbers were manually quantified in raw images using Fiji’s Multi-point tool. The center-to-center distance between thick filaments was measured following the methodology detailed by (Chakravorty et al., 2017). Average Z-disc images were created using TEM images of longitudinal DLM sections. Regions containing Z-bodies or Z-discs were then selected and compiled into a single stack. The stacks were aligned laterally with subpixel registration using the “Align slices in stack…” function of the Template Matching plugin (Tseng et al., 2012) in Fiji, and average projections were subsequently generated.

#### dSTORM images

The filtered and drift-corrected single-molecule localization data were processed as detailed in Szikora et al., 2020a, Szikora et al., 2020b, Varga et al., 2023. Briefly, visualization and quantitative analysis were performed using IFM Analyzer v2.1. To create averaged structures, the software’s merge function was used to align localizations along the symmetry axes of the H-zone/I-band before averaging. A super-resolved image with a 10 nm pixel size was then generated from the merged event lists. Relying on the symmetry axes of the H-zone and I-band, we aligned independently averaged protein densities to generate pseudo-multicolor representations. Quantitative measurements were conducted on averaged structures derived from either a single myofibril or, in cases of high noise, from multiple myofibrils captured within a single experiment.

### Simulated IFM Images

Ground truth images of phalloidin and α-Actinin labeled myofibrils were generated as 2D projections of uniformly labeled cylindrical objects, with overlapping regions representing Z-discs and gaps representing H-zones. These images were scaled and the image formation process was simulated in Fiji (Schindelin et al., 2012). Uneven labeling was modeled by applying a Gaussian noise (sigma = 2). The point spread function (PSF) of the microscope was simulated by convolving the images with a 2D Bessel function (numerical aperture: 1.4, wavelength: 525 nm) using Fiji’s MosaicSuite. Shot noise was introduced by adding Poisson noise.

### Data Analysis and Figures

Data collection and organization were carried out in Microsoft Excel, while statistical analysis and graph creation were performed using GraphPad Prism 8 or R. The raw data used to generate all plots presented in the figure panels are available in the corresponding source data files. Data normality was assessed using the D’Agostino & Pearson or the Shapiro-Wilk test. The specific statistical tests applied are detailed in the figure legends. Figures were created and finalized in Adobe Illustrator, and 3D models and animations were produced using Blender 4.0 (Blender Foundation).

## Supporting information

Fig1SourceData

Fig2SourceData

Fig3SourceData

Fig4SourceData

SFig1SourceData

SFig3SourceData

SFig5SourceData

## ACKNOWLEDGEMENTS

We thank Belinda Bullard, Anton Bryantsev, Velia Fowler, Florence Janody, the Developmental Studies Hybridoma Bank and the Bloomington Drosophila Stock Center for antibodies and fly stocks. We thank Elvira Czvik, Anikó Berente, and Erika Bánfiné Rácz for technical assistance.

---

**Supplementary Figure 1.**
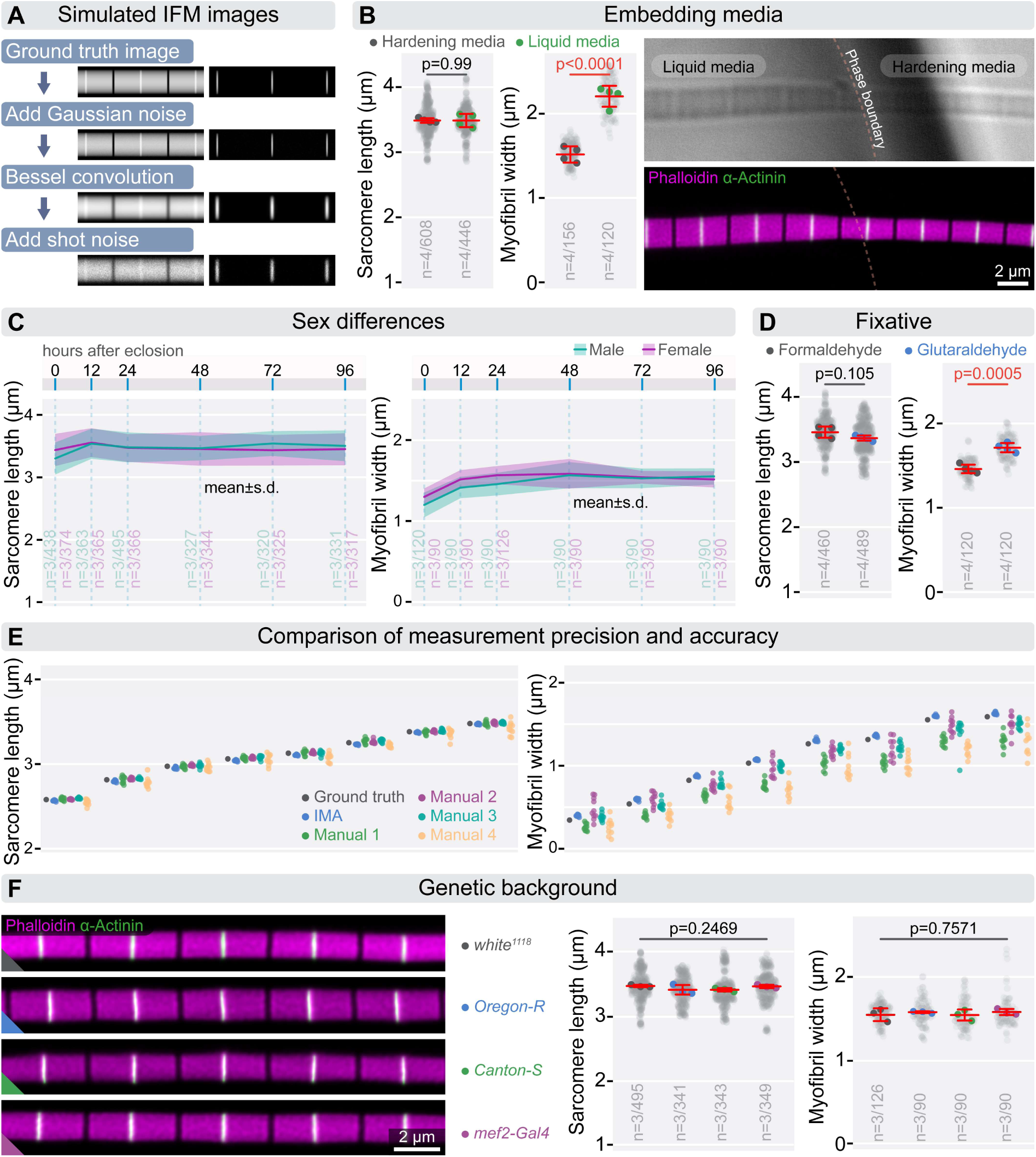
Morphometric measurements of IFM sarcomeres. **(A)** This panel illustrates how simulated IFM images with predefined diameters and sarcomere lengths were generated to compare the accuracy of different measurement methods (refer to the Methods section for details). **(B)** Panels highlight how embedding media affect sarcomere length and myofibril diameter. While sarcomere length remains unchanged (p=0.99), myofibril diameter is highly sensitive to the medium used, showing significant differences (p<0.0001). Hardening media, such as ProlongGold, reduce myofibril diameter, whereas liquid media, like glycerol-based solutions, increase it. Micrographs (on the right) illustrate this effect: in the transmitted light image (top), a liquid media bubble is visible within the hardening medium, and in the fluorescence image (bottom), variations in myofibril diameter are apparent. F-actin is stained with phalloidin (magenta), while Z-discs are labeled with α-Actinin (green). Statistical analysis was performed using an unpaired t-test. **(C)** Panels display the measured differences in sarcomere length and myofibril width between sexes after eclosion. **(D)** Panels show the effects of formaldehyde and glutaraldehyde on sarcomere length and myofibril diameter. While fixative choice does not significantly impact sarcomere length (p=0.105), myofibrils fixed with glutaraldehyde are significantly thicker than those fixed with formaldehyde (p=0.0005). An unpaired t-test was used for analysis. **(E)** These panels demonstrate the reliability of morphometric measurements performed using the Individual Myofibrils Analyzer (IMA) compared to manual measurements by four individuals on simulated images with predefined sarcomere length and diameter (dark gray dots). The simulation set included 80 images with varying sarcomere lengths and widths (10 images per category), randomly rotated. **(F)** The micrographs display representative images of isolated myofibrils from flies with different genetic backgrounds, labeled with phalloidin (magenta) and α-Actinin (green). The accompanying plots indicate no significant differences in the IFM morphometrics of these myofibrils. Statistical analysis was performed using one-way ANOVA with Tukey’s multiple comparison test. Light gray dots represent individual measurements of sarcomere length and myofibril diameter, while the larger dots indicate the mean values from independent experiments. Error bars represent the mean and s.d. of independent experiments. “n” refers to the number of independent experiments and the number of replicates. Raw data used to generate the plots presented in this figure are available in the source data file (SourceData).

**Supplementary Figure 2.**
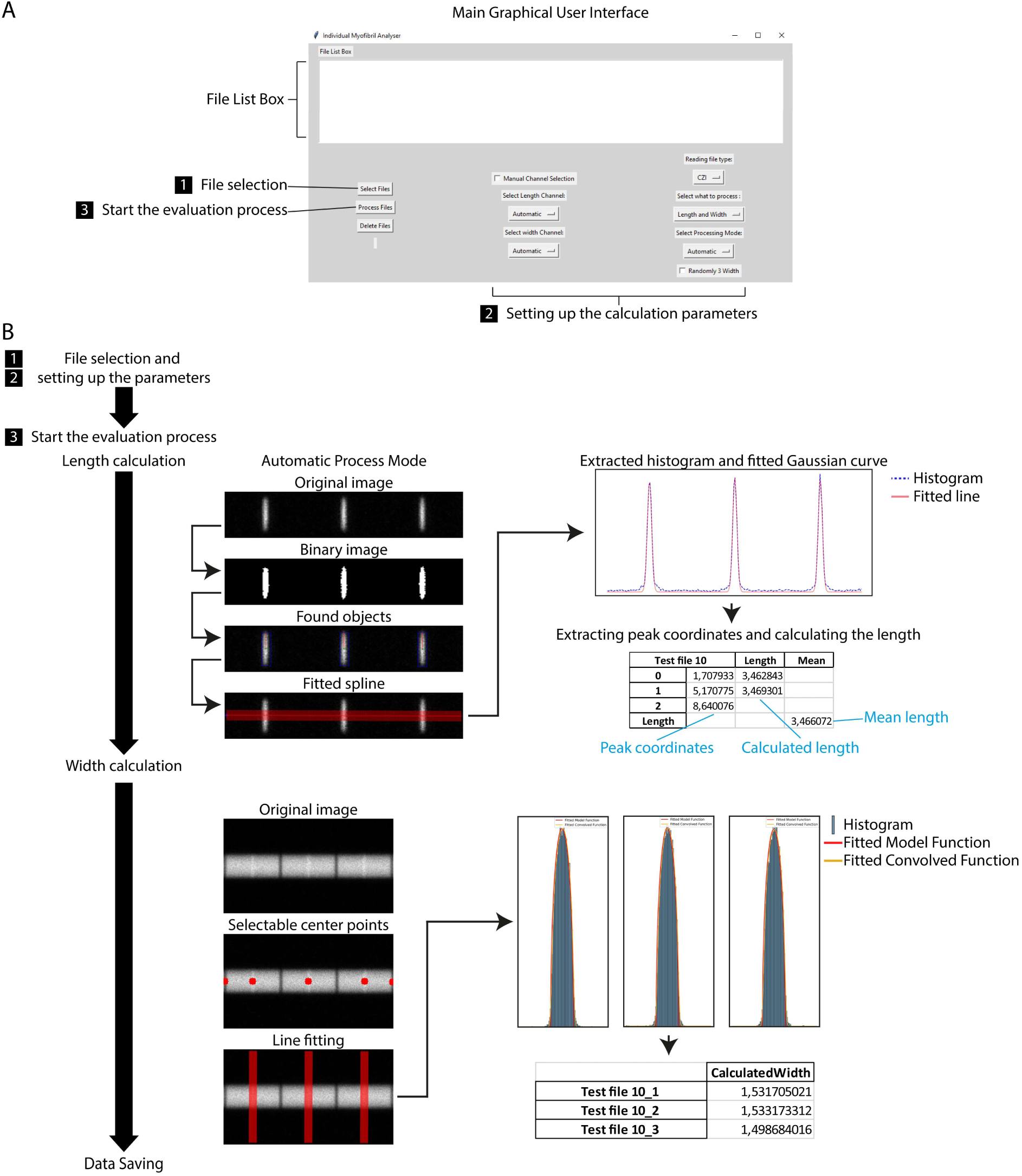
Key features and workflow of the Individual Myofibril Analyzer (IMA) **(A)** Graphical User Interface (GUI): The IMA’s user interface is designed for intuitive and efficient analysis. The interface is structured to facilitate a straightforward analysis with steps labeled from 1 to 3 guiding users through initiating evaluations within the software. **(B)** Analysis Workflow in Automatic Mode: After selecting the input file and configuring parameters, the software separates the image into two channels: “length” and “width.” In the “length” channel, a binary image is generated to identify objects. A spline connects the centers of the identified objects, and histogram data is extracted along the spline using 10-pixel averaging. The software then fits a Gaussian function to each peak, calculating the distances between them. For width measurement, the central points derived from the length analysis are used to draw perpendicular lines. Histogram data is extracted along these lines (again using 10-pixel averaging), and both a theoretical disk model and a convolved function are fitted to the data. This approach ensures precise estimation of myofibril width by accounting for ideal distributions and measurement-induced effects. The software saves the resulting histograms and calculated data as an Excel file. A comprehensive explanation of the software’s features and functions can be found in the User Guide (https://github.com/GorogPeter94/Individual-Myofibril-Analyser-IMA-/tree/main).

**Supplementary Figure 3.**
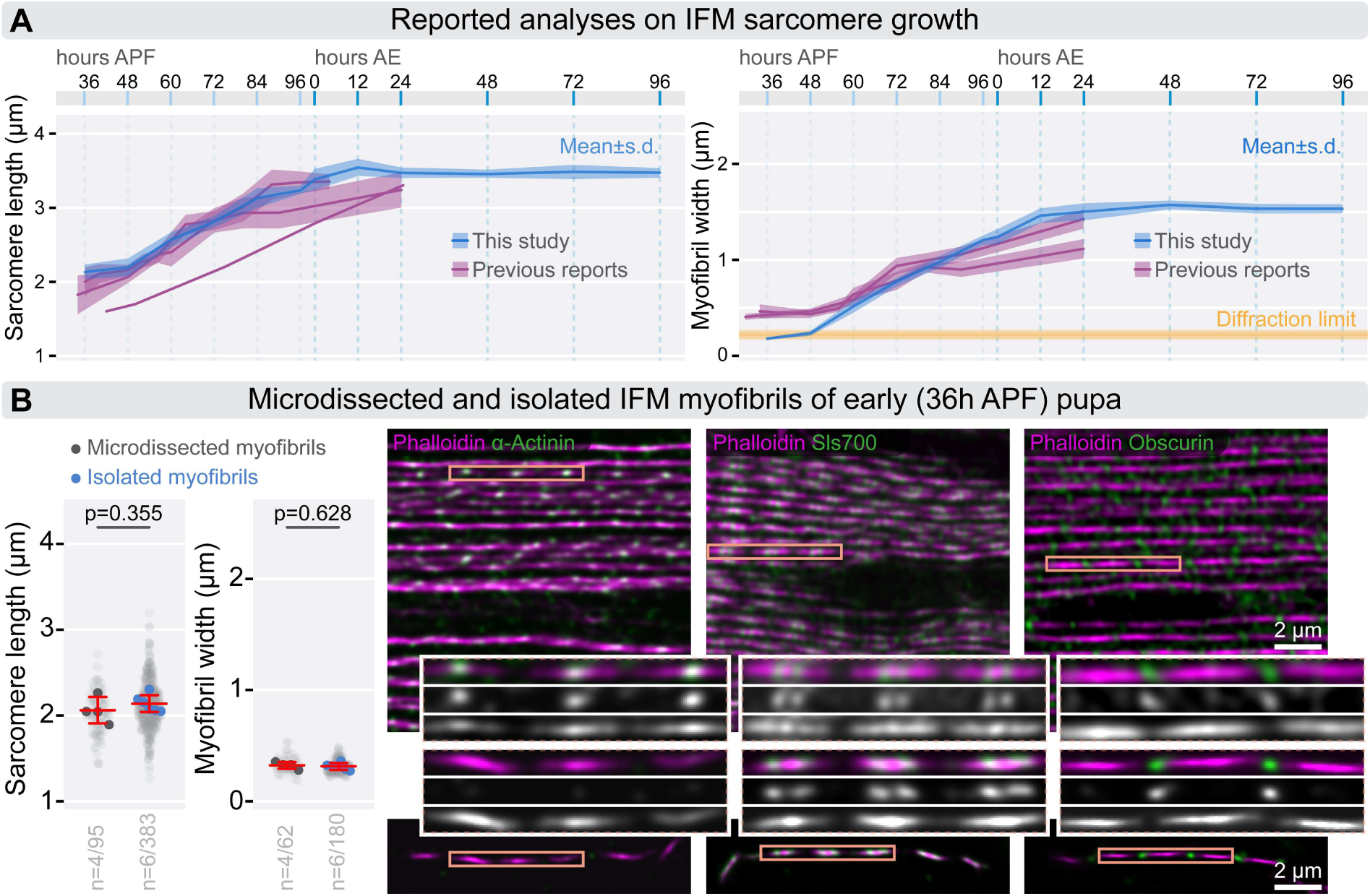
Comparison of IFM growth. **(A)** The plots compare previously published datasets on sarcomere length and myofibril width with the data obtained in this study. **(B)** Plots demonstrating that sarcomere length and myofibril diameter from young pupae (36 hours APF) are consistent whether measured in isolated individual myofibrils or within microdissected muscles. Light gray dots represent individual measurements, while larger dots show mean values from independent experiments. Error bars indicate the mean and s.d. across experiments, with “n” denoting the number of independent experiments and the number of replicates. The micrographs on the right compare the overall organization of microdissected muscles with isolated individual myofibrils from young pupae (36 hours APF). These images also display the localization of sarcomeric markers, including α-Actinin, Sls700 and Obscurin. Insets provide higher-resolution views to facilitate thorough comparisons. Raw data used to generate the plots presented in this figure are available in the source data file (SFig3SourceData).

**Supplementary Figure 4.**
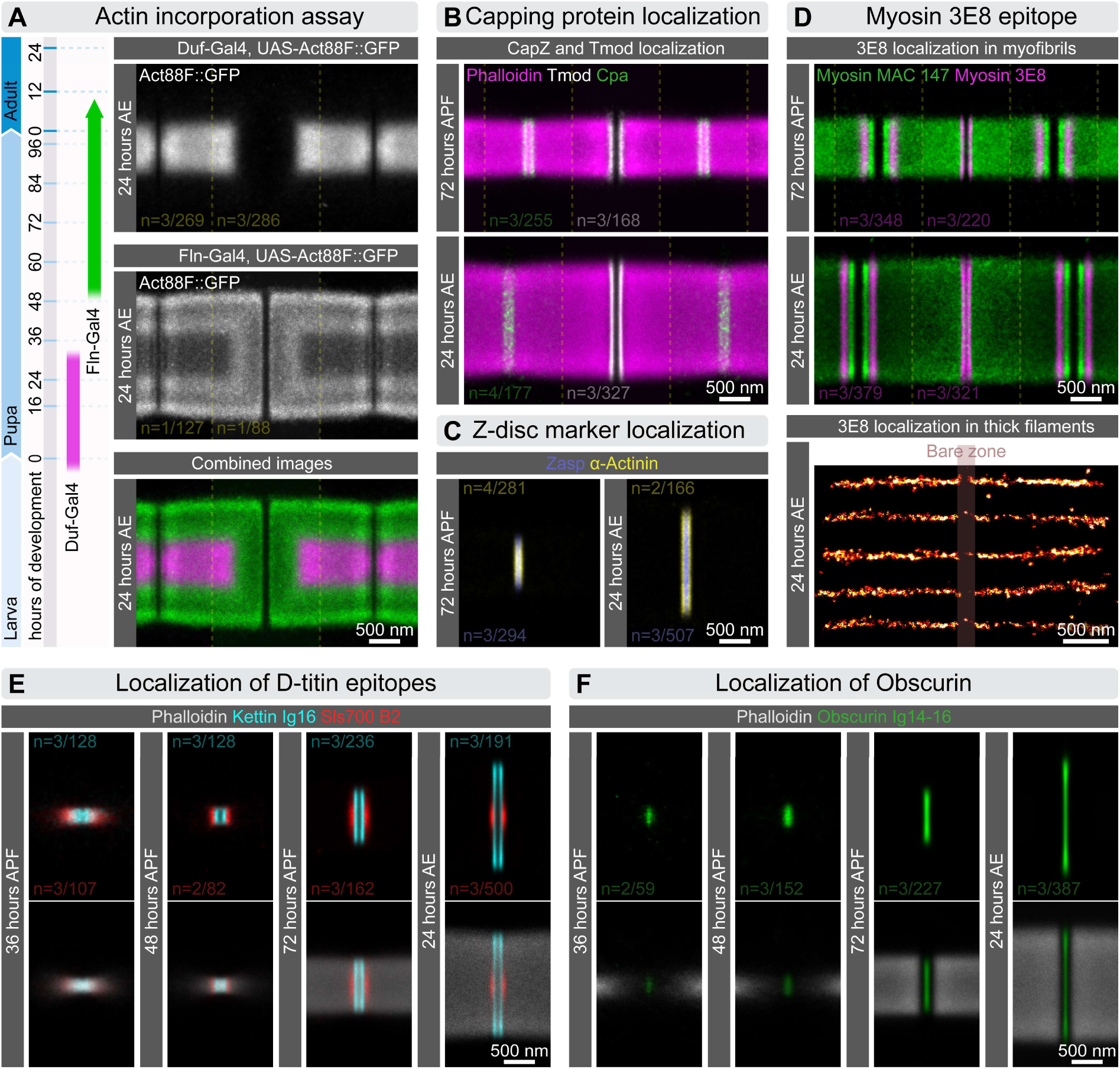
Nanoscopic localization of sarcomeric markers during myofibrillogenesis. **(A)** The schematic on the left illustrates the restricted temporal activation of the *duf-Gal4* and *fln-Gal4* drivers during myofibrillogenesis. On the right, dSTORM reconstructions display the incorporation of GFP-labeled Act88F monomers into IFM myofibrils using these drivers. **(B)** Pseudo-multicolor dSTORM reconstructions reveal the localization of Tmod (in white) and Cpa (in green) in IFM myofibrils isolated from pupae (72 hours APF) and adult flies (24 hours AE). F-actin is labeled with phalloidin (magenta). **(C)** Pseudo-multicolor dSTORM reconstructions highlight the localization of α-Actinin (yellow) and Zasp52 (blue) within the Z-disc of IFM myofibrils from pupae (72 hours APF) and adult flies (24 hours AE). **(D)** The top panels show multicolor dSTORM reconstructions of myosin epitopes (MAC 147 in green and 3E8 in magenta) in IFM myofibrils from pupae (72 hours APF) and adult flies (24 hours AE). At the bottom, dSTORM images show single isolated thick filaments from adult flies (24 hours AE) labeled with the 3E8 myosin antibody. **(E)** A series of pseudo-multicolor dSTORM reconstructions reveal the localization of elastic filament epitopes, Kettin Ig16 (cyan) and Sls700 B2 (red), at four different time points during myofibrillogenesis. F-actin is labeled with phalloidin (gray). **(F)** Series of dSTORM reconstructions illustrate the localization of the central Obscurin Ig14-16 (green) epitope, across four time points of myofibrillogenesis. F-actin is labeled with phalloidin (gray).

**Supplementary Figure 5.**
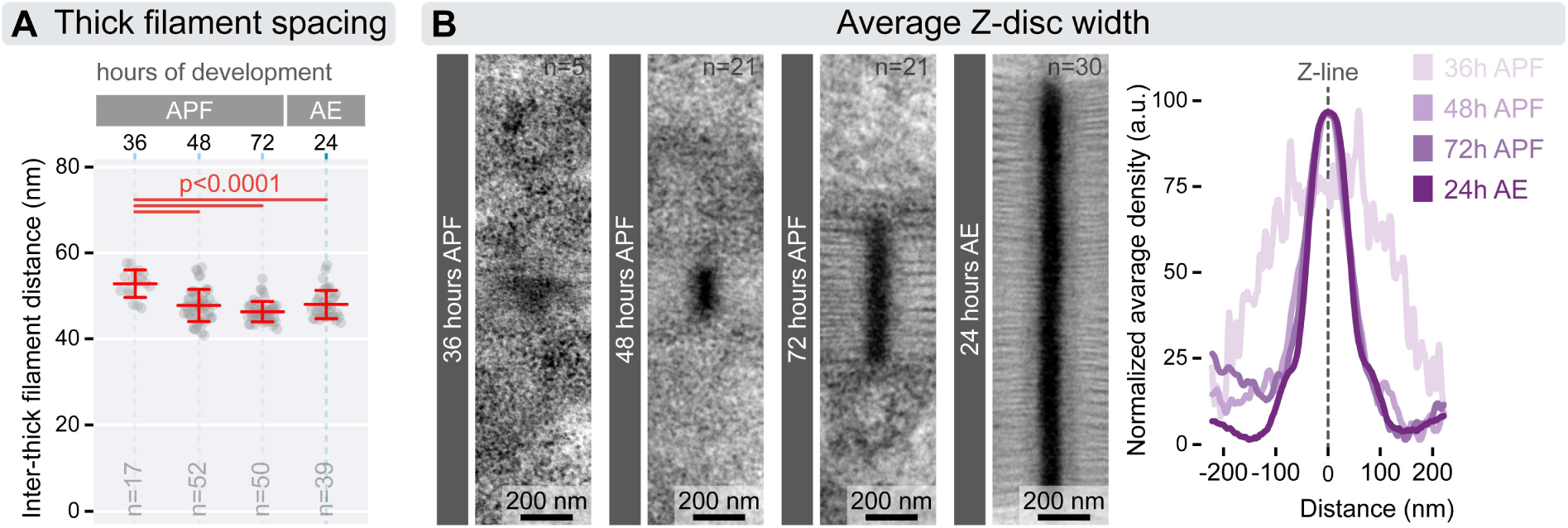
Myofilament lattice spacing and Z-disc width during myofibrillogenesis. **(A)** The plot shows the center-to-center spacing between thick filaments at four time points during myofibrillogenesis. At 36 hours APF, the spacing is around 53 nm, which significantly decreases to approximately 46-48 nm by 48 hours APF, and this spacing remains constant for the rest of development. Statistical analysis was performed using one-way ANOVA with Tukey’s multiple comparison test. Error bars represent the mean and s.d., and “n” refers to the number of myofibrils analyzed. **(B)** Averaged TEM images on the left illustrate the Z-body/disc morphology at four designated time points during myofibrillogenesis. The plot on the right shows changes in the Z-body/disc density profile throughout the process. At 36 hours APF, only diffuse Z-bodies are visible, which then organize into compact Z-discs by 48 hours APF. While the height of the Z-disc increases significantly, Z-disc width decreases only slightly from this point onward. Raw data used to generate the plots presented in this figure are available in the source data file (SFig5SourceData).

**Supplementary Figure 6.**
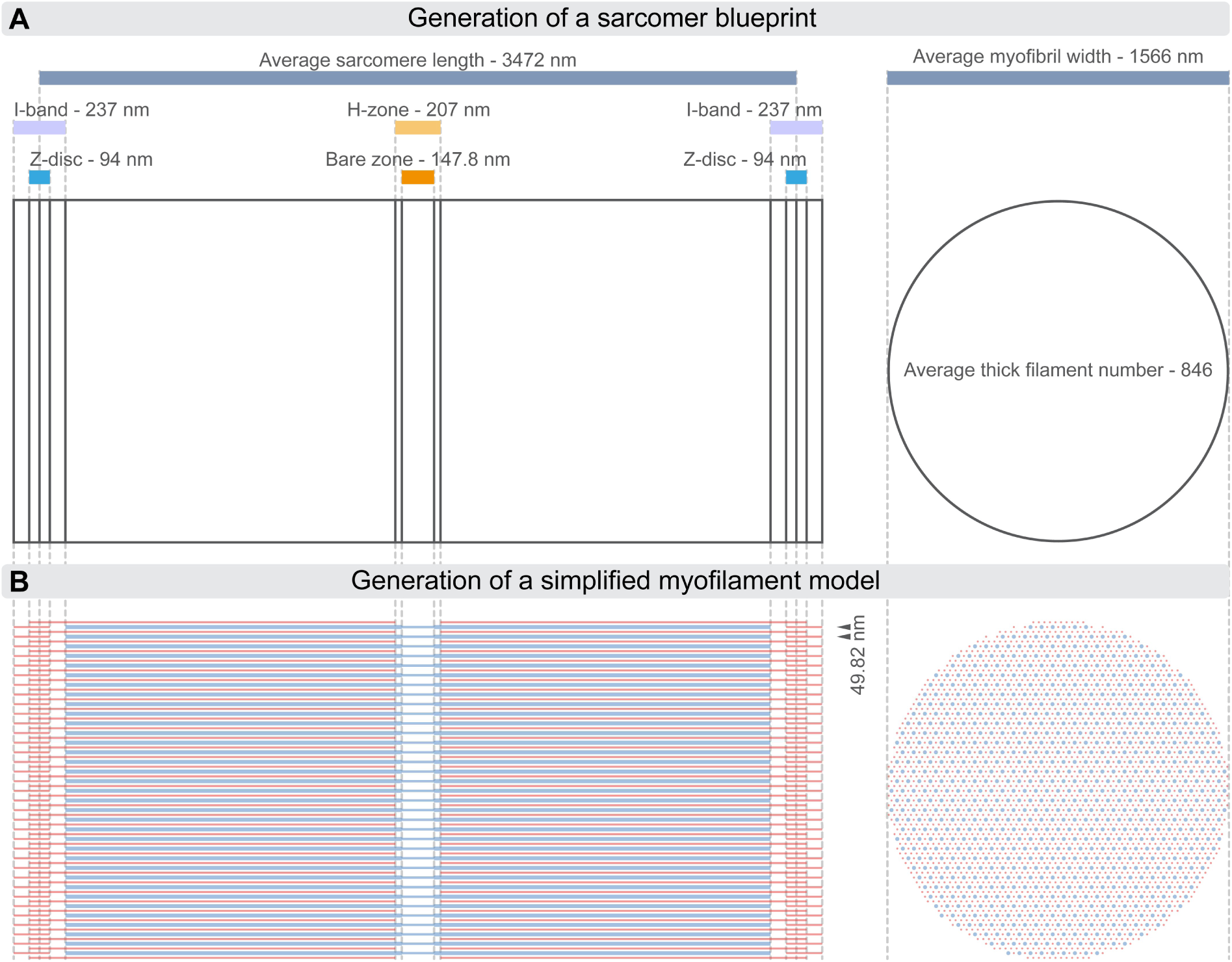
Outlining the ‘myofilament models’ of IFM sarcomeres. **(A)** The schematic outlines the process of creating a blueprint for a sarcomere model. The average length and diameter of the sarcomere are measured using confocal microscopy. The width of the H-zone, bare zone, I-band and Z-disc are measured from dSTORM images of phalloidin- or myosin-labeled myofibrils. The number of thick filaments is determined from TEM cross-sections of myofibrils. **(B)** Using these measurements, the myofilaments can be positioned within the model. To generate a cross-section, additional constraints are applied: Myofilaments arrange in a hexagonal pattern with a thick-to-thin filament ratio of 1:3. The filaments are organized into a MyAc layer, with thin filaments always positioned between two thick filaments. The center-to-center distance between thick filaments is then calculated, which is used to build the longitudinal model. This model allows the calculation of myofilament lengths: 1. Thick Filament Length (*L*_thick_): 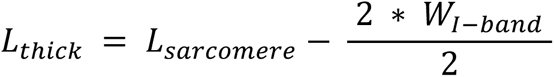 Where: *L*_*sarcomere*_ is the average sarcomere length *W*_*I-ba*_ is the average width of the I-band 2. Thin Filament Length (*L*_*thin*_): 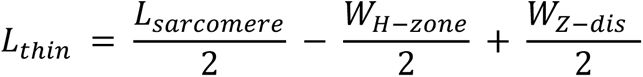 Where: *L*_*sarcomere*_ is the average sarcomere length *W*_*H-zon*_ is the average width of the H-zone *W*_*Z-disc*_ is the average width of the Z-disc

## Supplementary Movies 1-4

These animations illustrate the 3D lattice structure of an average sarcomere at specific time points during myofibrillogenesis: 36, 48 and 72 hours APF, as well as at 24 hours AE. Thick filaments are represented in blue, and thin filaments are shown in red. Each animation starts with a cross-sectional view of the sarcomere, which rotates 90° along the x-axis to reveal a side view. Then, the perspective shifts into an orthogonal view.

